# Adrenergic signaling controls early transcriptional programs during CD8^+^ T cell responses to viral infection

**DOI:** 10.1101/2021.10.20.465178

**Authors:** Leonardo D. Estrada, Didem Ağaç Çobanoğlu, Aaron Wise, Robert W. Maples, Murat Can Çobanoğlu, J. David Farrar

## Abstract

Viral infections drive the expansion and differentiation of responding CD8^+^ T cells into variegated populations of cytolytic effector and memory cells. While pro-inflammatory cytokines and cell surface immune receptors play a key role in guiding T cell responses to infection, T cells are also markedly influenced by neurotransmitters. Norepinephrine is a key sympathetic neurotransmitter, which acts to suppress CD8 ^+^ T cell cytokine secretion and lytic activity by signaling through the β2-adrenergic receptor (ADRB2). Although ADRB2 signaling is considered generally immunosuppressive, its role in regulating differentiation of effector T cells in response to infection has not been investigated. Using an adoptive transfer approach, we compared the expansion and differentiation of wild type (WT) to *Adrb2*^*-/-*^ CD8 ^+^ T cells throughout the primary response to vesicular stomatitis virus (VSV) infection *in vivo*. We measured the dynamic changes in transcriptome profiles of antigen-specific CD8 ^+^ T cells as they responded to VSV. Within the first 7 days of infection, WT cells out-paced the expansion of *Adrb2*^*-/-*^ cells, which correlated with reduced expression of IL-2 and the IL-2Rα in the absence of ADRB2. RNASeq analysis identified over 300 differentially expressed genes that were both temporally regulated following infection and selectively regulated in WT vs *Adrb2*^*-/-*^ cells. These genes contributed to major transcriptional pathways including cytokine receptor activation, signaling in cancer, immune deficiency, and neurotransmitter pathways. By parsing genes within groups that were either induced or repressed over time in response to infection, we identified three main branches of genes that were differentially regulated by the ADRB2. These gene sets were predicted to be regulated by specific transcription factors involved in effector T cell development, such as *Tbx21* and *Eomes*. Collectively, these data demonstrate a significant role for ADRB2 signaling in regulating key transcriptional pathways during CD8 ^+^ T cells responses to infection that may dramatically impact their functional capabilities and downstream memory cell development.

## Introduction

Cytolytic CD8^+^ T cells play a critical role in immune responses to pathogens and can be harnessed to target cancer. Their activation and development into effector cells are guided by a variety of signals that include antigen recognition, co-stimulatory receptor activation, and soluble factors such as cytokines. Upon antigen recognition, CD8^+^ T cells rapidly divide and acquire critical effector functions including cytokine secretion and lytic activity that are essential for pathogen clearance. Once the infection resolves, most cells die through attrition leaving a small pool of diverse memory cells with the capacity for rapid expansion and effector function in the face of a secondary infection. Many cells of the immune system, including CD8^+^ T cells, express various neurotransmitter receptors (1, 2), yet the role of neural signals in T cell function remains largely unexplored. As secondary lymphoid tissues are heavily innervated by post-ganglionic sympathetic neurons that secrete norepinephrine (NE), the sympathetic nervous system would be expected to play a pivotal role in immune regulation (2-4).

In previous studies, depletion of endogenous NE through chemical sympathectomy significantly enhanced the innate cytokine storm leading to exacerbated pathology during influenza infection of mice (5). This hyper-inflammation was accompanied by significantly increased IFN-γ-producing CD8^+^ T cells during the primary infection phase, indicating an important role for NE in limiting the magnitude of both innate and adaptive T cell responses to viral infections. More recent studies have demonstrated an intrinsic role for NE signaling to suppress cytokine secretion and lytic activity in both mouse and human CD8^+^ T cells (6-8). NE acted specifically and exclusively through the ADRB2 to modulate acute effector function. Thus, ADRB2 signaling plays a distinct role in limiting the magnitude of T cell-mediated primary responses.

Initial T cell activation pathways, such as antigen recognition and cytokine signaling, are critical to antiviral responses. We wished to understand how ADRB2 signaling impacted these early primary transcriptional responses of T cells to an *in vivo* virus infection. In this study, we assessed and compared detailed transcriptome changes of wild type (WT) and *Adrb2*-deficient (*Adrb2*^*-/-*^) CD8^+^ T cells throughout their responses to a viral infection and utilized novel bioinformatic tools to define regulatory elements controlled by adrenergic signaling. We found that intrinsic ADRB2 signaling in CD8^+^ T cells controls early transcriptional programs at all timepoints through the first 12 days of their response to Vesicular Stomatitis Virus (VSV) infection. Many of these dysregulated pathways belonged to known regulators of T cell function and development including cytokine signaling and response to pathogens. This study highlights a critical role for ADRB2 signaling in regulating dynamic transcriptome expression throughout CD8^+^ T cell antiviral responses.

## Results

To gain a better understanding of how ADRB2 signaling modulates peripheral CD8^+^ T cell development, we compared the expansion and gene expression changes that occurred in response to a virus challenge between WT and *Adrb2*^-/-^ CD8^+^ T cells. We utilized the clone4 T cell receptor transgenic (C4-Tg) model, which is specific for the influenza hemagglutinin antigen (HA) presented by H-2K^d^ on the BALB/c background (9). We measured antigen-specific T cell responses to a recombinant VSV expressing HA protein from influenza (VSV-HA) (10). In order to distinguish the responses, congenic WT (CD90.1/1) and *Adrb2*^-/-^ (CD90.1/2) C4-Tg T cells were adoptively co-transferred into BALB/c recipients (CD90.2/2), which then were infected with VSV-HA (Fig. 1A). Primary expansion was monitored in spleen and lymph nodes by staining for both CD90.1 and CD90.2 congenic markers that distinguished the transferred from the endogenous pool of CD8^+^ T cells as well as WT from *Adrb2*^-/-^ cells (Fig. 1B). WT and *Adrb2*^-/-^ cells expanded equally to day 5 following infection. However, we observed a significant reduction in *Adrb2*^-/-^ CD8^+^ T cells at days 7 and 12 post-infection, compared to WT, which correlated with reduced expression of the high-affinity IL-2Rα (CD25) on day 5 and lower IL-2 secretion from *in vitro*-stimulated cells (Supplemental Figs. 1A and B). Although we previously found no marked differences in either proliferation or apoptosis in response to antigen stimulation (11), we found that *Adrb2*^*-/-*^ cells displayed reduced induction of CD25 as a function of TCR-stimulated cell division *in vitro*, and supplementation of cultures with IL-2 restored CD25 expression in *Adrb2*^-/-^ cells (Supplemental Fig. 1C). In contrast, we found no significant differences in either *in vitro* cytokine secretion (IFN-γ and TNF-α) or lytic activity between WT and *Adrb2*^-/-^ cells at these early time points of infection (data not shown). Thus, the attenuated early proliferation of *Adrb2*^-/-^ cells correlated with their reduced IL-2 and IL-2Rα expression, yet their cytokine secretion potential on a per-cell basis remained intact during their expansion into effector cells.

**Figure 1.**
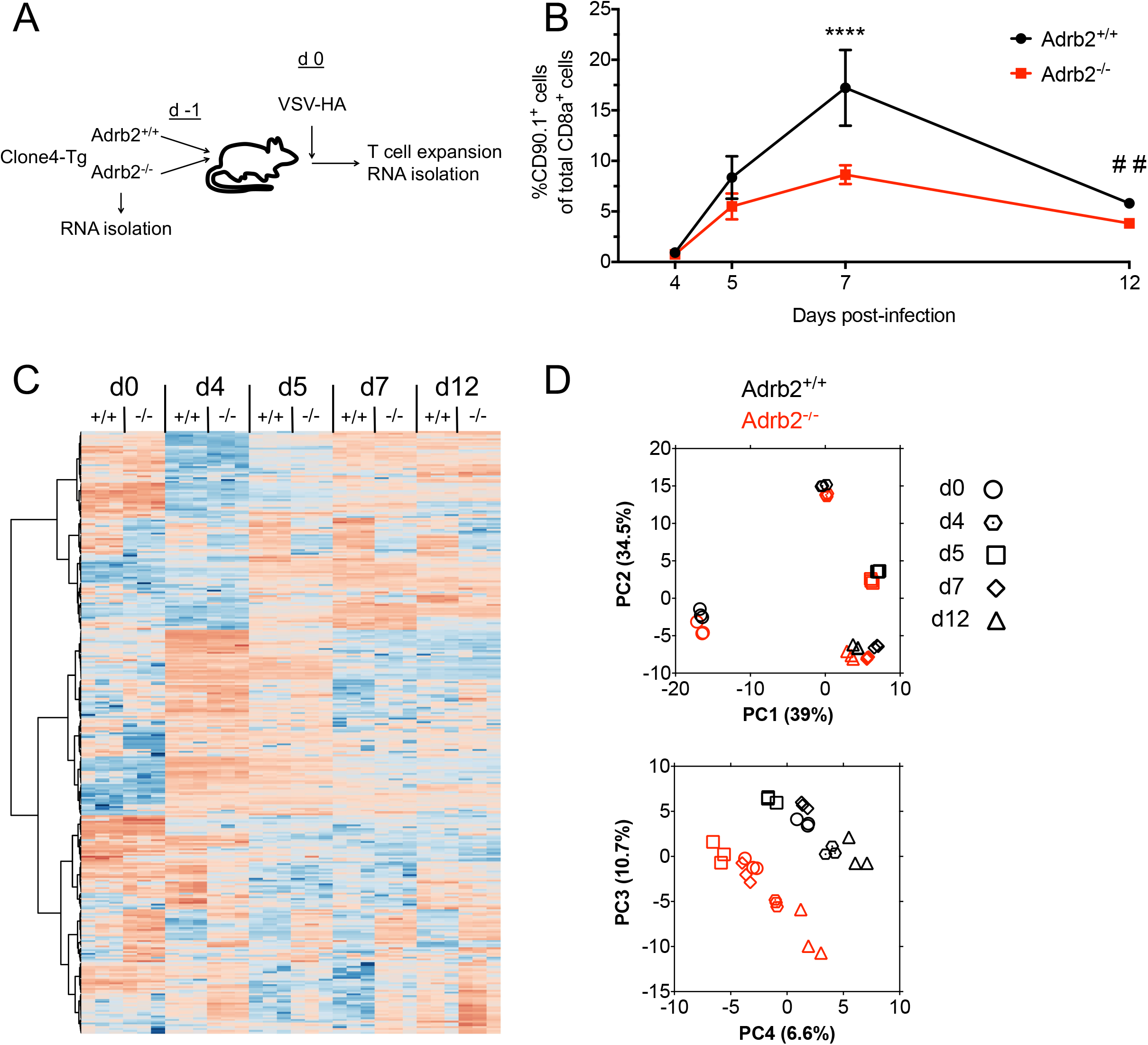
The ADRB2 regulates transcriptional programs during early T cell priming. (**A**) WT (CD90.1/1) and *Adrb2*^-/-^ (CD90.1/2) C4-Tg T cells were co-transferred (1:1, 1000 cells each) to BALB/cJ (CD90.2/2) recipients, which were infected with VSV-HA. RNA was isolated for RNASeq analysis from purified cells on day 0 and from infected hosts on days 4, 5, 7, and 12 post-infection. (**B**) Expansion of transferred cells in pooled spleen and lymph node was quantified by flow cytometry by measuring the percentage of WT and *Adrb2*^-/-^ cells within the total proportion of congenic CD90.1^+^ cells (**** p < 0.0001 and ## p <0.01 by two-way ANOVA). (**C**) EdgeR analysis identified all genes differentially expressed at any timepoint between WT and *Adrb2*^-/-^ cells. Gene expression values were used to perform 1-way hierarchical clustering, and data are displayed as a heat map. (**D**) Principal component analysis of differentially expressed genes between WT and *Adrb2*^-/-^ cells displayed as a function of each timepoint post-infection (PC1 vs PC2, top panel; PC3 vs PC4, bottom panel).

Early T cell priming events regulate long-range transcriptional programs that lead to both effector and memory cell development (12, 13). We measured gene expression changes in FACS-purified WT and *Adrb2*^-/-^ cells from VSV-HA-infected animals at each time point shown in Fig. 1B by RNASeq analysis. Over 6000 genes were collectively regulated in response to infection in both cell types. EdgeR analysis (14) identified over 320 genes that were differentially expressed between WT and *Adrb2*^-/-^ cells at any of the time points, including day 0 (Supplemental Table 1). A one-way hierarchical cluster of these genes demonstrated a temporal change in gene expression, which for some clusters of genes differed at all timepoints (Fig. 1C). The temporal dynamics of gene expression changes were highlighted by principal component (PC) analysis of this gene set, as genes within the first two PCs differed significantly by time, but not by genotype (Fig. 1D, upper panel). These components were driven by the expression of genes involved in the primary effector response such as cytokines, chemokines, and cytokine receptors. However, PCs 3 and 4 displayed a marked division over time between WT and *Adrb2*^-/-^ cells, which were comprised of genes involved in a variety of cellular processes including transcription, signal transduction, and cellular differentiation (Fig. 1D, lower panel). We further segregated these temporal gene sets based on their up or down-regulation at each time point relative to WT cells, and select genes are annotated within the volcano plots in Fig. 2A. Genes that were significantly differentially expressed, either positively or negatively, between WT and *Adrb2*^-/-^ cells were then assessed for their contribution to specific KEGG pathways (Fig. 2B). We found that unique pathways were engaged by ADRB2 signaling at incremental times through their progression to effector cells. Among these pathways, we found that select pathways were dysregulated at multiple times throughout the early phase of infection, such as cytokine receptor interaction, transcriptional misregulation in cancer, and circadian rhythms. The genes driving these varied pathways are listed in Supplemental Table 2 and included *Stat1, Il2ra, Il10ra, Per2, Fbxl3*, and *Mef2c*. Although the sorted populations of cells were determined to be >95% pure in post sorting analyses, we observed low-level expression of some B cell-associated mRNAs including *Cd19* and *Btk*, which were differentially expressed in *Adrb2*^*-/-*^ cells. These data suggest an important role for the ADRB2 in regulating the temporal expression of genes during the early stages of T cell priming.

**Figure 2.**
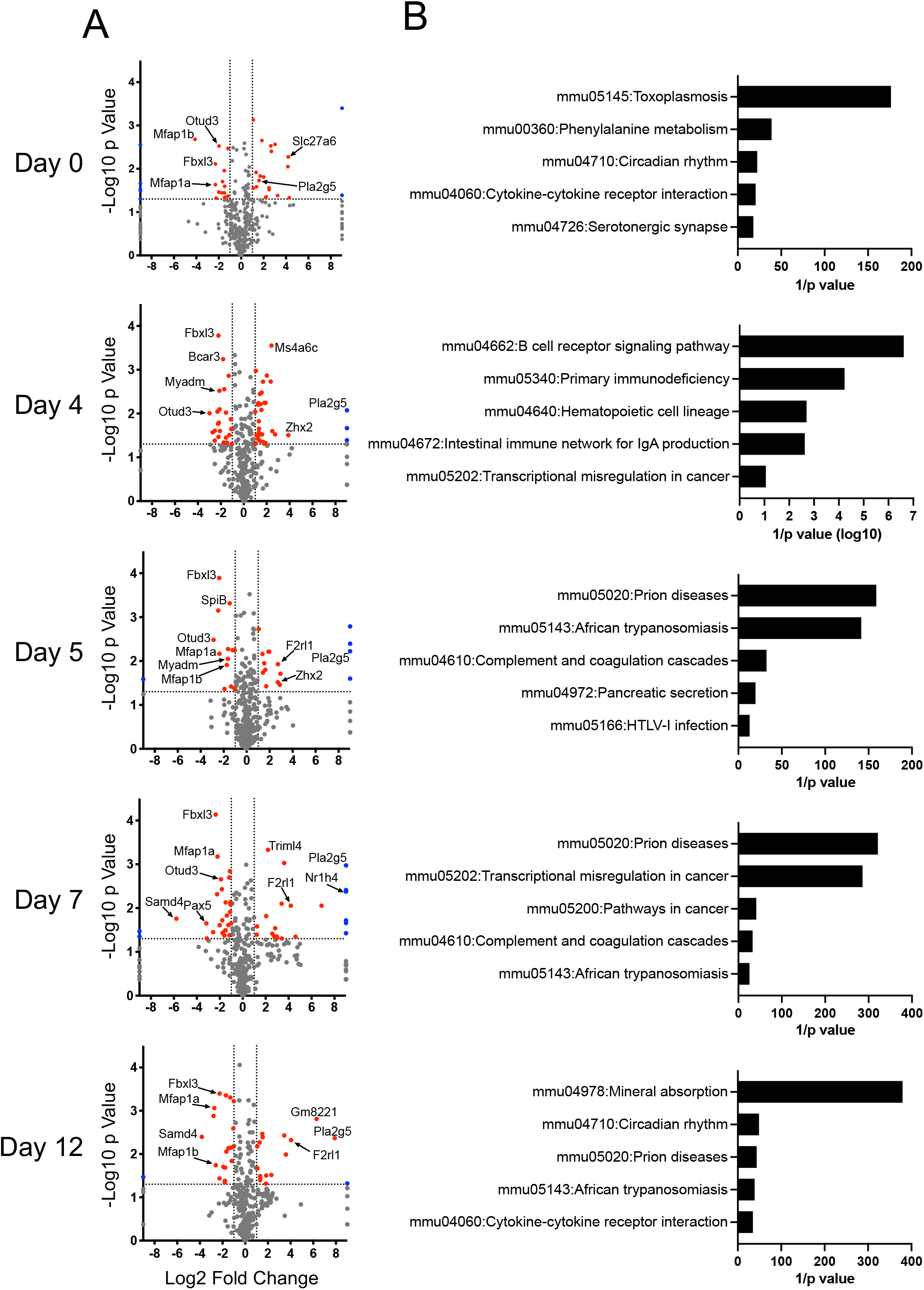
The ADRB2 selectively regulates transcriptional pathways involved in diverse immune functions. Differentially expressed genes between WT and *Adrb2*^-/-^ cells at each time point of infection are displayed in volcano plots, and select genes contributing to major KEGG pathways are denoted within the plots. (B) KEGG pathway analysis was performed with the differentially expressed genes at each time point. The top five pathways are displayed.

As cells divide and develop over time, their gene expression patterns become highly variegated. These fluctuations in gene expression can be modeled in terms of the regulatory dynamics that cause them. We utilized SMARTS to model, compare, and visualize the regulatory dynamics following infection for WT and *Adrb*2^-/-^ cells (Fig. 3A) (15). Using SMARTS, we constructed regulatory models for each condition using CD8^+^-specific transcription regulator (TR)-gene interaction data derived from Best et al. (16). Genes were parsed into paths based on the similarity they share in both the direction and magnitude of expression at each time point, as well as the TRs which are known to regulate them. A full list of genes for each path is provided in Supplemental Tables 3-4. Each SMARTS model represents sets of genes following a similar regulatory program as paths; split nodes represent regulatory events that cause groups of genes to diverge in expression. Select TRs for each path are listed in box diagrams for each path in Fig. 3A, and a complete list is provided in Supplemental Tables 5-6. As expected, genes within both positively and negatively regulated paths shared regulation by select transcription factors known to be involved in CD8^+^ T cell function and memory development including *Tbx21, Eomes, Irf5, Rxra, Prdm1, Id2*, and *Stat4*. Gene regulation by these TRs was predicted by SMARTS to be shared between WT and *Adrb*2^-/-^ cells. SMARTS was also used to identify TRs that follow distinct regulatory programs in WT and *Adrb*2^-/-^ cells (Supplemental Table 7). TRs found to be involved in differential gene regulation within select paths are highlighted in red in Fig. 3A. For example, *Mbd2* was predicted to selectively regulate genes in WT paths C and H, while *Mafb* was predicted to regulate genes within path E in *Adrb*2^-/-^ cells but not WT cells.

**Figure 3.**
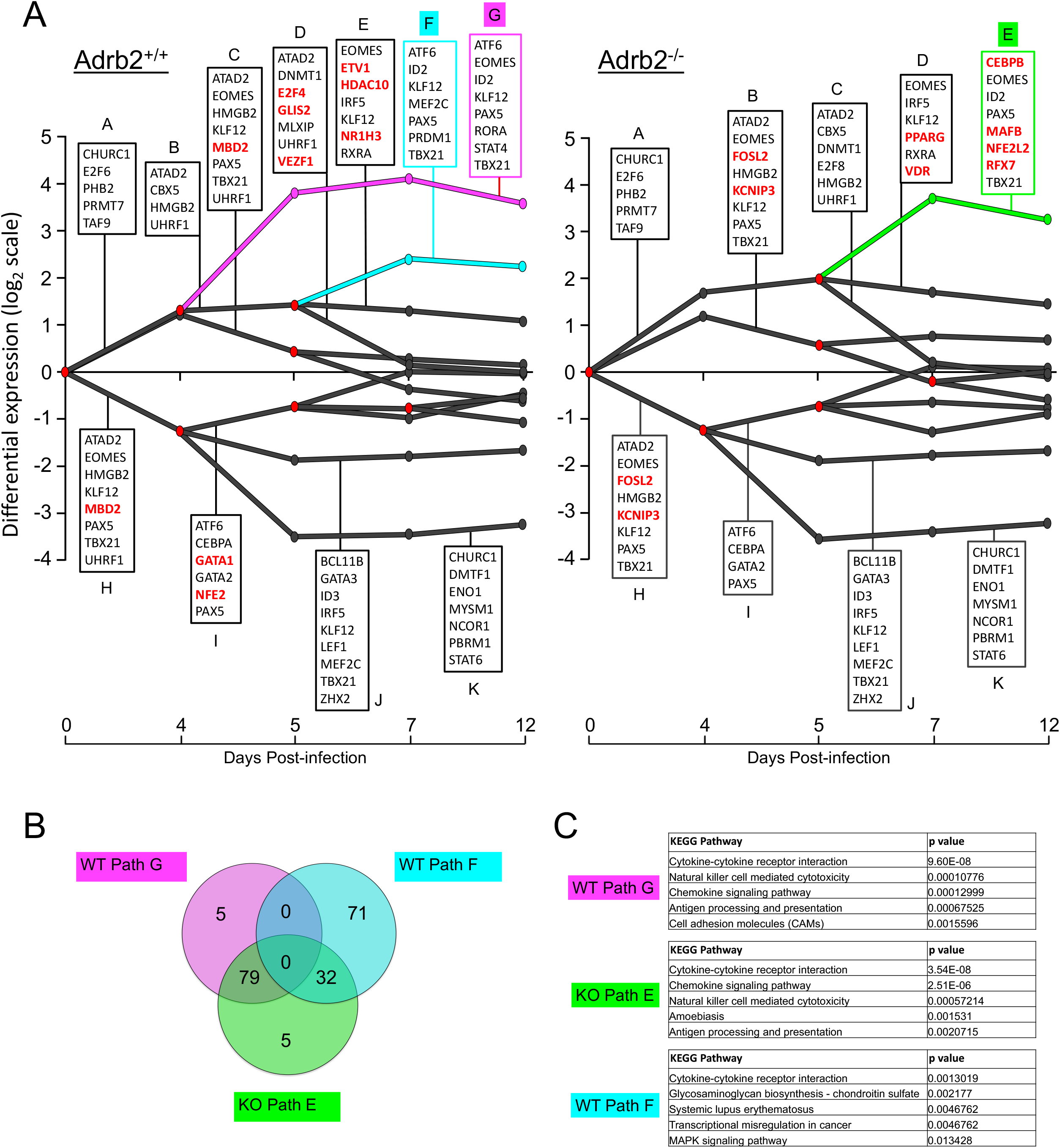
ADRB2 signaling coordinates temporal waves of gene expression. (**A**) SMARTS analysis at each timepoint compared to the day 0 pre-transfer condition. Each path corresponds to clusters of genes sharing common magnitude, direction of expression and regulatory factors. Red nodes in the model represent splits in expression between groups of genes. TRs predicted to regulate each split path are listed above and below their corresponding paths, and TRs in red were predicted by SMARTS to be involved in differential gene regulation between WT and *Adrb2*^-/-^ cells. WT paths F (cyan) and G (magenta), and *Adrb2*^-/-^ path E (green) contained genes and regulators which were significantly altered between the two models; they were subjected to further analyses (in B and C). (**B**) Venn analysis of constituent genes within WT paths F and G and *Adrb2*^-/-^ path E. Values represent the numbers of genes within each unique or shared segment. (**C**) Integrated KEGG pathway analysis of constituent genes of WT paths F and G and *Adrb2*^-/-^ path E. The top 5 pathways from each path are listed.

Overall, the overall pattern of gene expression changes that occurred over time were similar between WT and *Adrb*2^-/-^ cells. However, we identified a unique path consisting of genes that were more highly induced on day 4 post-infection in WT cells that were not regulated at that timepoint in *Adrb*2^-/-^ cells (Fig. 3A, *Adrb2*^+/+^ path G (magenta)). By comparing the constituent genes of paths F (cyan) and G (magenta) in WT cells with path E (green) in *Adrb*2^-/-^ cells, the majority of genes in WT path G were included in the split path E in *Adrb*2^-/-^ cells (Fig. 3B, Supplemental Table 8). Further, the genes in WT path G and *Adrb*2^-/-^ path E mapped to many of the top KEGG pathways involved in T cell effector function (Fig. 3C). Despite their similar directional change, only 1/3 of genes within WT path F were shared with *Adrb*2^-/-^ path E, which was underscored by the unique KEGG pathways involved in WT path F. These data suggest that the effector response genes induced at day 4 post-infection in WT cells were either temporally delayed or absent in their induction in *Adrb*2^-/-^ cells.

Common to both paths in WT and *Adrb*2^-/-^ cells, *Tbx21* and *Eomes* were predicted by SMARTS to regulate genes known to be involved in the transition from effector to memory cells, yet *Tbx21* itself was not differentially expressed at any time point between the two models (Fig. 4A). However, *Eomes* was more highly expressed in *Adrb*2^-/-^ than WT cells on days 7 and 12 post-infection (Fig. 4B), which may impact the effector to memory transition based on previous studies. Within the gene sets of WT path G and *Adrb*2^-/-^ path E, we identified several TRs that were differentially expressed at day 4 post-infection, the time point at which the major split path occurred in WT but not *Adrb*2^-/-^ cells. These factors included *Prdm1, Pax5, Spib, Mef2c, Mafb*, and *Bach2* (Fig. 4C-H). Of these factors, *Pax5* and *Spib* are predominantly expressed in B cells and silenced in T cells (17, 18), yet their transient induction in WT CD8^+^ T cells during infection may indicate a previously unanticipated role for them in T cell function. However, *Prdm1, Mef2c*, and *Bach2* have been shown to regulate various aspects of T cell function (16, 19, 20), specifically the effector to memory cell transition. Of note, *Prdm1* was not only more highly induced in WT compared to *Adrb*2^-/-^ cells at day 4 post-infection, it was also selectively included in genes constituent to WT path G, but not in *Adrb*2^-/-^ path E (Supplemental Table 6). Thus, the ADRB2 is involved in regulating the proper timing of gene expression patterns during early T cell priming, and the factors that correlated with these differences are known regulators of effector T cell development.

**Figure 4.**
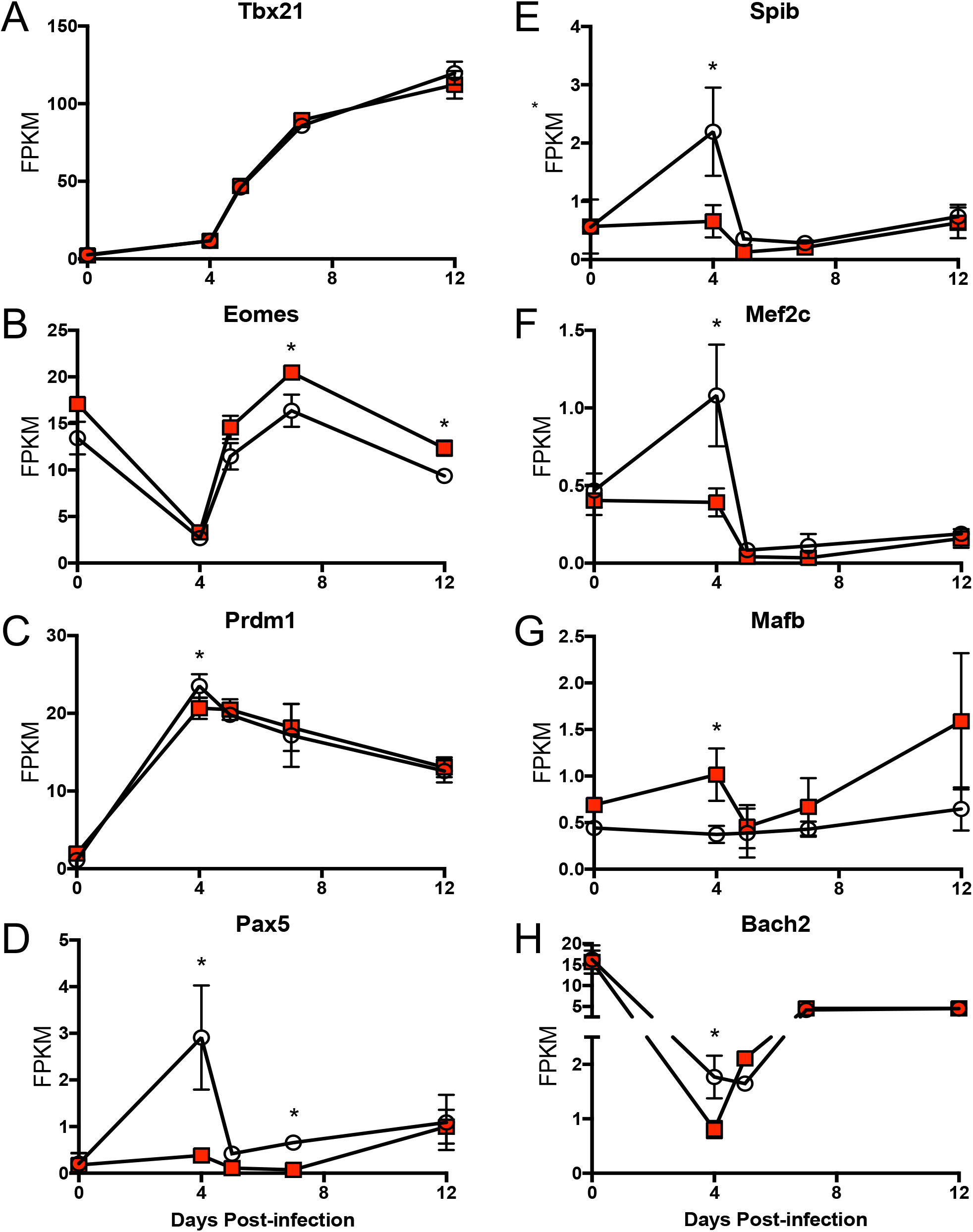
Select transcription factor expression regulated by the ADRB2. Normalized FPKM values for specific transcription factors are displayed for WT (open circles) and *Adrb2*^*-/-*^ (red squares) CD8^+^ T cells isolated at the indicated time points of infection. (A) *Tbx21*, (B) *Eomes*, (C) *Prdm1*, (D) *Pax5*, (E) *Spib*, (F) *Mef2c*, (G) *Mafb*, (H) *Bach2*.

## Discussion

The sympathetic nervous system controls a broad range of behavioral and physiological processes. Given the significant innervation of secondary lymphoid tissues by sympathetic neurons, it is not surprising that this neurotransmitter pathway regulates immune function. Indeed, prior reports have shown a clear role for NE and the ADRB2 in suppressing a variety of acute immune functions (21), including acute cytokine secretion and lytic activity in CD8^+^ T cells (5-8). In prior studies, the absence of adrenergic signaling *in vivo*, by chemical sympathectomy, increased IFN-γ producing CD8^+^ T cells after influenza infection (5). Consistent with these findings, we recently demonstrated that antigen-induced cytokine secretion from CD8^+^ T cells was markedly decreased when mice were treated with a long-acting β2-agonist during VSV infection (6). Although NE and ADRB2 agonists suppressed acute CD8+ T cell effector function in response to antigen receptor activation, it was not clear how adrenergic signaling influenced downstream effector T cell development. Our previous studies found that *in vitro* priming of naive CD8^+^ T cells into effector cells was not affected by NE during the differentiation stage (6). In the current study, we uncovered a key role for ADRB2 signaling that regulated various aspects of CD8^+^ T effector cell responses to *in vivo* virus infection. First, we found that adrenergic signaling regulated CD25 expression and IL-2 secretion, which correlated with a reduced proportion of *Adrb2*^*-/-*^ to WT Ag-specific cells towards the end of the expansion phase *in vivo*. As CD25 expression correlates with memory cell development (22-24), it is possible that the transition from effector to memory cell development can be impacted by ADRB2 signaling. Second, our current work demonstrated that intrinsic ADRB2 signaling on CD8^+^ T cells regulates a dynamic program of gene expression that correlates with both their expansion and their time-dependent development into effector cells.

Gene expression is regulated by networks of interactions, and these networks show context-dependent adaptation (25, 26). The dynamic changes in gene expression that occurred in CD8^+^ T cells during their expansion into effector cells was generally preserved in *Adrb2*^*-/-*^ cells, as the main components of those gene clusters evolved over time in both WT and *Adrb2*^*-/-*^ cells. However, by monitoring the dynamic changes in gene expression that occurred throughout the effector expansion phase with SMARTS (15), we identified clusters of genes that were significantly dysregulated in the absence of the *Adrb2*. Further analyses predicted unique transcription factors that could be responsible for the regulation of those gene clusters based on combinatorial gene expression (27). The main branch of genes that were induced in WT cells at d4 post-infection (path G) shared most of its genes with those induced in *Adrb2*^*-/-*^ cells at a later time (d5, path E). While most of the predicted regulators of those genes were common to both paths, some were more highly induced at d4 in WT cells, such as *Pax5* and *Mafb*. The d4 path G in WT cells was comprised of a variety of genes involved in major T cell fate-determining pathways, and the delay in their induction observed in *Adrb2*^*-/-*^ cells may indicate a critical role for *Adrb2* signaling in the temporal response to infection. It is unlikely that ADRB2 signaling regulates a single factor that solely orchestrates these branch points. Nonetheless, we speculate that the *Adrb2* regulated those factors during T cell proliferation to provide a temporal program of gene expression corresponding to their timely response to the pathogen. These mechanisms could include signaling pathways, transcriptional changes, post-translational modifications, epigenetic alterations, and asymmetric inheritance of fate-determining factors during cell division (23, 28, 29).

Previous studies have demonstrated an immunosuppressive role for ADRB2 signaling, acting acutely to dampen the magnitude of cytokine expression and lytic activity in pre-committed effector cells (8, 30, 31). Interestingly, deletion of *Adrb2* also impacts the diurnal recirculation of T cells through secondary lymphoid tissues (32, 33), indicating a role for ADRB2 signaling in circadian regulation. Indeed, we identified several core circadian genes, such as *Per2* and *Fbxl3*, differentially regulated at various time points of infection, suggesting a direct role for regulating these clock genes intrinsically in CD8^+^ T cells. The detailed temporal map of gene expression described here has identified a clear role for the ADRB2 in the primary response of acute CD8^+^ T effector cells to viral infection. Future studies will utilize these networks to identify how these pathways functionally regulate effector and memory cell development through the course of infection.

## Acknowledgments

Primary RNASeq data were deposited in the GEO database under accession number GSE102478 (“Timeseries analysis of gene expression in Adrb2+/+ and Adrb2-/-C4-transgenic CD8+ T cells responding to Vesicular Stomatitis Virus expressing hemagglutinin from influenza A PR/8”).

The authors thank Dr. Virginia Sanders (Ohio State University) for sharing the BALB/c x *Adrb2*^-/-^ animals. We thank Ms. Angela Mobley and the UT Southwestern flow cytometry facility for excellent cell sorting assistance. We are grateful to Drs. Lora Hooper, Regina Rowe, and Michelle Gill for their advice on this project and for reviewing the manuscript.

This work was supported by NIH grants AI056222, AI125545, and AI143248 (JDF). LDE was supported by NIH training grant AI005284. DA was supported by a Careers in Immunology fellowship from the American Association of Immunologists.

## Author contributions

LDE designed and performed experiments and wrote the paper; DA performed experiments and edited the paper; AW performed SMARTS analysis and wrote the paper; MCC designed bioinformatic approach; and JDF designed the study, performed data analysis, and wrote the paper.

## Materials and Methods

### Animals

BALB/cJ, Clone4-Tg (Cl4) (34), and *Adrb2*^-/-^ (35) mice were housed in specific pathogen-free conditions at the University of Texas Southwestern Medical Center Animal Research Center facilities. Adrb2^-/-^ mice bred onto the BALB/c background (36) were a kind gift from Dr. Virginia Sanders (Ohio State University), and Cl4 mice were purchased from Jax mice (Jackson laboratory). All experiments involving mice in this study were approved by the Institutional Animal Care and Use Committee of the University of Texas Southwestern Medical Center.

### Adoptive T cell transfer and VSV-HA infection

Recombinant VSV-HA expressing hemaglutinin from Influenza A PR/8 (10) was a kind gift from Dr. Elizabeth Ramsburg. CD8^+^ T cells were isolated from spleen and lymph nodes (axillary, brachial, inguinal, and superficial cervical) of WT and *Adrb2*^-/-^ Cl4 mice (7-12 weeks old) with a negative isolation kit according to the manufacturer’s instructions (Invitrogen #11417D). Donor T cells were derived from CD90.1 congenic animals in order to track their frequency in CD90.2 hosts. For co-transfer experiments, 2000 cells of a 1:1 mix of each genotype (1000 cells each) were intravenously (i.v.) injected into naive BALB/cJ mice in 100 µLs of sterile saline. One day later, each mouse was infected i.v. with 1e^6^ plaque-forming units (PFU) of VSV-HA. Mice were then sacrificed at the time points indicated post infection, and the frequency of transferred cells was monitored by flow cytometry by staining for CD90.1 and CD90.2.

### In vitro T cell assays

*In vitro* cytokine assays were performed as previously described (6). Single cell suspensions from spleen and lymph nodes (axillary, brachial, inguinal and superficial cervical) were prepared separately and incubated at 2×10^6^/mL in the presence or absence of the CD8^+^ T cell-specific HA peptide (IYSTVASSL, 50 nM) for 21-24 hrs. For *in vitro* cell division assays, cells were pre-labeled with CFSE prior to activation and allowed to divide in culture for 72 hrs. Cells were stained for CD90.1 and CD25 and analyzed by FACS.

### RNASeq Analysis

WT and *Adrb2*^-/-^ C4-Tg T cells were co-transferred to BALB/cJ recipients followed by infection with VSV-HA, as described above. RNA was isolated from cells prior to transfer and from CD90.1/2 FACS-purified cells isolated from cohorts of infected animals at incremental days post-infection. WT and *Adrb2*^-/-^ cells were distinguished based on single or co-expression of CD90.1 and CD90.2. Barcoded libraries were prepared from purified mRNA (New England Biolabs, #E7530S, #E7490S, #E7335S, and Axygen #MAG-PCR-CL-5) and sequenced on an Illumina HISEQ 2500.

Quality assessment of the RNASeq data was performed using NGS-QC-Toolkit (37). Reads with more than 30% of nucleotides with Phred quality scores less than 20 were removed from further analysis. Quality filtered reads were then aligned to the mouse reference genome GRCm38 (mm10) using the HISAT (v 2.0.1) aligner (38) using default setting except for – library-type = fr-firststrand. Aligned reads were counted using featureCount (v1.4.6) (39) per gene ID. Differential gene expression analysis was performed using the R package edgeR (14) (v 3.8.6). For each comparison, genes were required to have 1 read in at least 1 sample to be considered as expressed. They were used for normalization factor calculation. Gene differential expression analysis was performed using GLM approach following edgeR analysis. Cutoff values of fold change greater than 2 and FDR less than 0.01 were then used to select for differentially expressed genes between sample group comparisons. Normalized gene FPKM values were averaged within groups for heatmap generation. MetaboAnalyst3.0 (40) was used to perform R-based principle component analysis (PCA) and integrated pathway analysis using the KEGG metabolic pathway database.

To perform the SMARTS analysis, we developed a new version of SMARTS, ‘Supervised SMARTS’ that can use known class labels for building models. SMARTS uses an Input/Output Hidden Markov Model (IOHMM) approach to model the regulation of genes over time. Each SMARTS model reconstructs the regulatory activity of an ensemble of individual time series. SMARTS requires as input a mapping between transcriptional regulators and their gene targets. We generated such a mapping using the regulatory behavior identified in Best et al. in their supplementary table 12 (16). We used SMARTS to build two models, one from the three WT cells time series and the other from the three *Adrb2*^-/-^ cells time series. The SMARTS analysis covered all 5 time points, from 0-12 days post infection. We further used SMARTS to identify putative differentially active transcription factors between the two models. In brief, we identify transcription factors whose regulated genes can only have their gene expression patterns explained by the proper model. This criterion is evaluated using a permutation test to determine statistical significance. See Wise and Bar-Joseph, 2013 section 2.4 for complete details (15).

### Statistical analyses

Three different statistical tests were performed using the GraphPad Prism software. For simple pairwise comparisons, a Student’s two-tailed t-test was used. Otherwise, a one-way or two-way ANOVA was used followed by a Bonferroni posttest for pairwise comparisons within the groups, as indicated in the figure legends. Differences were considered significant at p ≤ 0.05.

## Supplementary Materials

**Supplementary Figure 1: The *Adrb2* controls expression of IL-2 and the IL-2Rα**.

**Supplementary Tables S1-S8:**

**S1 – Table of all differentially expressed genes between WT and *Adrb2***^**-/-**^ **cells at all timepoints**.

**S2 – KEGG analysis of differentially expressed genes between WT and *Adrb2***^***-/-***^ **cells at each timepoint**.

**S3 – Tables of genes in each path of the *Adrb2***^**+/+**^ **SMARTS model**.

**S4 – Tables of genes in each path of the *Adrb2***^**-/-**^ **SMARTS model**.

**S5 – Tables of transcription factor predictions for each path of the *Adrb2***^**+/+**^ **SMARTS model**.

**S6 – Tables of transcription factor predictions for each path of the *Adrb2***^**-/-**^ **SMARTS model**.

**S7 – Table of transcriptional regulator (TR) prediction differences between *Adrb2***^**+/+**^ **and *Adrb2***^**-/-**^ **models**.

**S8 – Table of genes included in unique and shared paths by Venn analysis**.

## Notes

https://www.ncbi.nlm.nih.gov/geo/query/acc.cgi?acc=GSE102478

### Competing Interest Statement

The authors have declared no competing interest.

## References

1. D. M. Nance, V. M. Sanders, Autonomic innervation and regulation of the immune system (1987-2007). Brain. Behav. Immun. 21, 736–745 (2007).

2. D. Sharma, J. D. Farrar, Adrenergic regulation of immune cell function and inflammation. Semin. Immunopathol. 42, 709–717 (2020).

3. S. S. Chavan, K. J. Tracey, Essential Neuroscience in Immunology. J. Immunol. 198, 3389–3397 (2017).

4. D. Agac, M. A. Gill, J. D. Farrar, Adrenergic Signaling at the Interface of Allergic Asthma and Viral Infections. Front. Immunol. 9, 736 (2018).

5. K. M. Grebe et al., Sympathetic nervous system control of anti-influenza CD8+ T cell responses. Proc Natl Acad Sci U S A 106, 5300–5305 (2009).

6. L. D. Estrada, D. Agac, J. D. Farrar, Sympathetic neural signaling via the beta2-adrenergic receptor suppresses T-cell receptor-mediated human and mouse CD8(+) T-cell effector function. Eur J Immunol 46, 1948–1958 (2016).

7. A. Zalli et al., Targeting β2 adrenergic receptors regulate human T cell function directly and indirectly. Brain Behav Immun 45, 211–218 (2015).

8. C. Slota, A. Shi, G. Chen, M. Bevans, N. P. Weng, Norepinephrine preferentially modulates memory CD8 T cell function inducing inflammatory cytokine production and reducing proliferation in response to activation. Brain. Behav. Immun. 46, 168–179 (2015).

9. D. Lo et al., Peripheral tolerance to an islet cell-specific hemagglutinin transgene affects both CD4+ and CD8+ T cells. Eur. J. Immunol. 22, 1013–1022 (1992).

10. B. E. Barefoot, K. Athearn, C. J. Sample, E. A. Ramsburg, Intramuscular immunization with a vesicular stomatitis virus recombinant expressing the influenza hemagglutinin provides post-exposure protection against lethal influenza challenge. Vaccine 28, 79–89 (2009).

11. L. D. Estrada, D. Agac, J. D. Farrar, Sympathetic neural signaling via the beta2-adrenergic receptor suppresses T-cell receptor-mediated human and mouse CD8+ T-cell effector function. Eur. J. Immunol. 10.1002/eji.201646395 (2016).

12. J. Arsenio et al., Early specification of CD8+ T lymphocyte fates during adaptive immunity revealed by single-cell gene-expression analyses. Nat. Immunol. 15, 365–372 (2014).

13. F. Z. Chowdhury, H. J. Ramos, L. S. Davis, J. Forman, J. D. Farrar, IL-12 selectively programs effector pathways that are stably expressed in human CD8+ effector memory T cells in vivo. Blood 118, 3890–3900 (2011).

14. M. D. Robinson, D. J. McCarthy, G. K. Smyth, edgeR: a Bioconductor package for differential expression analysis of digital gene expression data. Bioinformatics 26, 139–140 (2010).

15. A. Wise, Z. Bar-Joseph, SMARTS: reconstructing disease response networks from multiple individuals using time series gene expression data. Bioinformatics 31, 1250–1257 (2015).

16. J. A. Best et al., Transcriptional insights into the CD8(+) T cell response to infection and memory T cell formation. Nat. Immunol. 14, 404–412 (2013).

17. A. Souabni, C. Cobaleda, M. Schebesta, M. Busslinger, Pax5 promotes B lymphopoiesis and blocks T cell development by repressing Notch1. Immunity 17, 781–793 (2002).

18. B. Bartholdy et al., The Ets factor Spi-B is a direct critical target of the coactivator OBF-1. Proc. Natl. Acad. Sci. U. S. A. 103, 11665–11670 (2006).

19. A. Kallies, A. Xin, G. T. Belz, S. L. Nutt, Blimp-1 transcription factor is required for the differentiation of effector CD8(+) T cells and memory responses. Immunity 31, 283–295 (2009).

20. M. J. Richer, M. L. Lang, N. S. Butler, T Cell Fates Zipped Up: How the Bach2 Basic Leucine Zipper Transcriptional Repressor Directs T Cell Differentiation and Function. J. Immunol. 197, 1009–1015 (2016).

21. D. Agac, L. D. Estrada, R. Maples, L. V. Hooper, J. D. Farrar, The beta2-adrenergic receptor controls inflammation by driving rapid IL-10 secretion. Brain. Behav. Immun. 74, 176–185 (2018).

22. K. C. Verbist et al., Metabolic maintenance of cell asymmetry following division in activated T lymphocytes. Nature 532, 389–393 (2016).

23. J. T. Chang et al., Asymmetric T lymphocyte division in the initiation of adaptive immune responses. Science 315, 1687–1691 (2007).

24. J. B. Spangler et al., Antibodies to Interleukin-2 Elicit Selective T Cell Subset Potentiation through Distinct Conformational Mechanisms. Immunity 42, 815–825 (2015).

25. K. Mitra, A. R. Carvunis, S. K. Ramesh, T. Ideker, Integrative approaches for finding modular structure in biological networks. Nat. Rev. Genet. 14, 719–732 (2013).

26. T. Ideker, N. J. Krogan, Differential network biology. Mol. Syst. Biol. 8, 565 (2012).

27. A. Wise, Z. Bar-Joseph, cDREM: inferring dynamic combinatorial gene regulation. J. Comput. Biol. 22, 324–333 (2015).

28. T. A. Doering et al., Network analysis reveals centrally connected genes and pathways involved in CD8+ T cell exhaustion versus memory. Immunity 37, 1130–1144 (2012).

29. C. D. Scharer, A. P. Bally, B. Gandham, J. M. Boss, Cutting Edge: Chromatin Accessibility Programs CD8 T Cell Memory. J. Immunol. 198, 2238–2243 (2017).

30. L. D. Estrada, D. Ağaç, J. D. Farrar, Sympathetic neural signaling via the β2-adrenergic receptor suppresses T-cell receptor-mediated human and mouse CD8(+) T-cell effector function. Eur. J. Immunol. 46, 1948–1958 (2016).

31. K. M. Grebe et al., Sympathetic nervous system control of anti-influenza CD8+ T cell responses. Proc. Natl. Acad. Sci. U. S. A. 106, 5300–5305 (2009).

32. A. Nakai, Y. Hayano, F. Furuta, M. Noda, K. Suzuki, Control of lymphocyte egress from lymph nodes through beta2-adrenergic receptors. J. Exp. Med. 211, 2583–2598 (2014).

33. K. Suzuki, Y. Hayano, A. Nakai, F. Furuta, M. Noda, Adrenergic control of the adaptive immune response by diurnal lymphocyte recirculation through lymph nodes. J. Exp. Med. 213, 2567–2574 (2016).

34. D. Lo et al., Peripheral tolerance to an islet cell-specific hemagglutinin transgene affects both CD4+ and CD8+ T cells. Eur J Immunol 22, 1013–1022 (1992).

35. D. K. Rohrer, A. Chruscinski, E. H. Schauble, D. Bernstein, B. K. Kobilka, Cardiovascular and metabolic alterations in mice lacking both beta1-and beta2-adrenergic receptors. J Biol Chem 274, 16701–16708 (1999).

36. J. W. McAlees, V. M. Sanders, Hematopoietic protein tyrosine phosphatase mediates beta2-adrenergic receptor-induced regulation of p38 mitogen-activated protein kinase in B lymphocytes. Mol Cell Biol 29, 675–686 (2009).

37. R. K. Patel, M. Jain, NGS QC Toolkit: a toolkit for quality control of next generation sequencing data. PLoS One 7, e30619 (2012).

38. D. Kim, B. Langmead, S. L. Salzberg, HISAT: a fast spliced aligner with low memory requirements. Nat. Methods 12, 357–360 (2015).

39. Y. Liao, G. K. Smyth, W. Shi, featureCounts: an efficient general purpose program for assigning sequence reads to genomic features. Bioinformatics 30, 923–930 (2014).

40. J. Xia, D. S. Wishart, Using MetaboAnalyst 3.0 for Comprehensive Metabolomics Data Analysis. Curr Protoc Bioinformatics 55, 14 10 11–14 10 91 (2016).

