## Supplemental Figure 1 for "Adrenergic signaling controls early transcriptional programs during CD8^+^ T cell responses to viral infection"

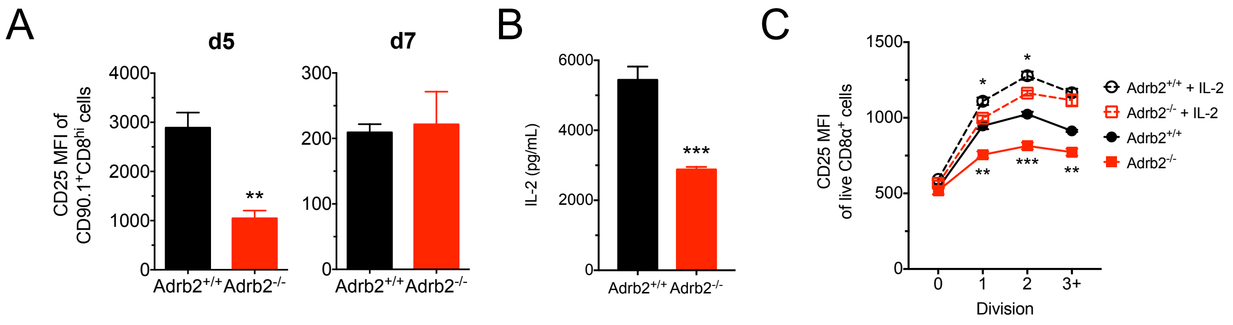


**Supplemental Figure 1**. The *Adrb2* controls expression of IL-2 and the IL-2Rα. (A) WT and *Adrb2*$^{-/-}$ cells from VSV-HA-infected animals were distinguished based on expression of CD90.1 and assessed for relative expression of the high affinity IL-2Rα (CD25) at days 5 and 7 after infection. (B) Spleen cell cultures from WT and *Adrb2*$^{-/-}$ CL-4 TCR Tg animals were cultured with HA peptide for 24 hrs. IL-2 was measure by ELISA from the culture supernatants. (C) WT and *Adrb2*$^{-/-}$ CL-4 TCR Tg spleen cells were labeled with CFSE and activated for 72 hrs with HA peptide in the absence or presence of IL-2 (50 U/ml). Relative expression of CD25 was measured in live/CD8+ cell by FACS.
